# Multiplex PCR for simultaneous genotyping of kdr mutations V410L, V1,016I and F1,534C in *Aedes aegypti* (L.)

**DOI:** 10.1101/2019.12.17.880195

**Authors:** Karina Villanueva-Segura, Gustavo Ponce-Garcia, Beatriz Lopez-Monroy, Esteban de J. Mora-Jasso, Lucia Perales, Francisco J. Gonzalez-Santillan, Kevin Ontiveros-Zapata, Jesus A. Davila-Barboza, Adriana E. Flores

## Abstract

**Background:** Knock down resistance (kdr) is the main mechanism that confers resistance to pyrethroids and DDT. This is a product of nonsynonymous mutations in the he voltage-gated sodium channel (VGSC) gene, these mutations produce a change of a single amino acid which reduces the affinity of the target site for the insecticide molecule. In Mexico, V410L, V1,016I and F1,534C mutations are common in pyrethroid-resistant *Aedes aegypti* (L.) populations.

**Methods:** A multiplex PCR was developed to detect the V410L, V1,016I and F1,534C mutations in *Ae. aegypti*. The validation of the technique was carried out using wild populations previously characterized for the three mutations through allele-specific PCR (AS-PCR) and with different levels of genotypic frequencies.

**Results:** The standardized protocol for multiplex endpoint PCR was highly effective in detecting 12 genotypes in five wild *Ae. aegypti* populations from Mexico. A complete concordance with AS-PCR was found for the simultaneous detection of the three kdr mutations.

**Conclusions:** Our diagnostic method is highly effective for the simultaneous detection of V410L, V1,016I and F1,534C, when they co-occur. This technique represents a viable alternative to complement and strengthen current monitoring and resistance management strategies against *Ae. aegypti*.

## Background

The portfolio of insecticides available for the control of arthropod pathogen vectors is very limited and is unlikely to increase in the near future, mainly due to the high cost of developing new molecules and products. Therefore, the emergence of resistance to commonly used insecticides is a serious threat to our ability to fight diseases transmitted by *Aedes aegypti* (L.). The development of such resistance is a complex and dynamic process dependent on many factors, so its management requires two types of information: good knowledge of the resistance mechanisms and their monitoring. The characterization of the resistance mechanisms involved allows us to assess and predict their impact on vector control strategies. Having routine monitoring of insecticide resistance in natural vector populations helps us to detect early resistance and improve the effectiveness of operational control strategies. *Ae. aegypti* is the main vector of dengue, chikungunya and Zika virus in Mexico, and its control depends largely on the use of pesticides that vary in their mode of action and include organophosphates, carbamates and neonicotinoids, but pyrethroids remain the preferred class, due to their fast action, high insecticidal activity and low toxicity to mammals [1, 2]. The target site of pyrethroids is the voltage-gated sodium channel (VGSC) present in the axon membrane of neurons and excitable cells in insects, and these insecticides produce a knockdown effect, that is, instantaneous paralysis in the insect due to prolonged activation and subsequent blockage of the action potentials of these channels [1, 3, 4, 5].

The main mechanisms that confer resistance to pyrethroids are overexpression of detoxifying enzymes and/or insensitivity at the target active site [6], being the mechanism associated with detoxification enzymes identified in *Ae. aegypti* populations from different regions of Mexico [7, 8, 9], particularly the mechanisms associated with esterases, glutathione S-transferases and mixed-function oxidases. On the other hand, knockdown resistance is conferred principally by nonsynonymous mutations that reduce pyrethroid binding to VGSC. In Mexico, it has been determined that pyrethroid resistance in *Ae. aegypti* is associated with high frequencies of any of the V1,016I, F1,534C and V410L mutations or combinations thereof. V1,016I was the first to be reported in a population of *Ae. aegypti* from Isla Mujeres, Quintana Roo, Mexico resistant to permethrin [10]. Subsequently, Ponce-García *et al*. demonstrated through a retrospective analysis carried out with 78 collections of *Ae. aegypti* that this replacement was practically absent in samples collected between 1996 and 2001, showing a dramatic increase between 2007 and 2009 [11]. Additionally, Siller et al. reported an increase in allele I1,016 frequency in *Ae. aegypti* populations in 2009 in several locations in Mexico [12]. Vera-Maloof *et al*. performed a linkage disequilibrium analysis in populations collected in Mexico between 2000 and 2012 that carried I1,016 and C1,534, and their results suggested that the sequential evolution of both mutations was necessary for pyrethroid resistance to develop [13]. Saavedra-Rodríguez *et al*. reported the V410L mutation in *Ae. aegypti* collections obtained during 2002 to 2016 from several locations in Mexico observing a high frequency of V410L in collections previously genotyped with V1,016I and F1,534C [14]. Later, Villanueva-Segura *et al*. reported high allelic frequencies of L410 in 25 populations of *Ae. aegypti* occurring in 2018 in eastern and southern Mexico with frequencies of 0.10-0.99 [15].

Currently, there are different PCR-based techniques for the detection of kdr mutations, which offer high sensitivity and specificity. However, when choosing one of these techniques, it is important to consider the economic resources of the laboratory, the training of technical personnel and the time available [16]. This point is even more important when considering the geographic extent that the vector occupies, the quantity of samples to be processed and the dynamics and fluctuation of the kdr resistance in short periods of time. The high-performance techniques available for genotyping kdr mutations, which are characterized by their speed and high sensitivity, are: sequencing of specific regions of VGSC, real-time PCR with TaqMan probes and high-resolution melting (HRM). However, given the need to acquire specialized equipment and supplies and the quantity of samples to be processed, the high cost of these methods constitutes its greatest disadvantage. The above has made AS-PCR one of the most used techniques for this purpose, offering a low cost and low error rate. Despite this, for laboratories in developing or low-budget countries, the use of other techniques based on allele-specific PCR is still recommended, when considering cost and ease of use and performance [17, 18. 19. 20]. Recently, the multiplex PCR technique has proven effective in the identification of two kdr mutations (V1,016G and F1,534C) simultaneously in *Ae*. aegypti [21]. In addition, it is possible to adapt it to different biochemical assays that not only reveal the amplified products optimally, but also contribute to improving the reliability, cost and performance of resistance monitoring [22].

The aim of this study was to develop a multiplex PCR method to detect three mutations, V410L, V1,016I and F1,534C, in *Ae. aegypti* in a single reaction. This technique can reduce the cost and time spent to monitor allelic frequencies in many countries where all three mutations co-occur.

## Materials and Methods

### Collection sites and laboratory conditions

Field collections of *Ae. aegypti* were carried out in 2018 from three states in Mexico: Nuevo Leon in the northeast with two locations, Monterrey and Guadalupe; Yucatan in the southeast with two locations, Merida and San Antonio Kaua; and Chiapas in the south with one location.

Mosquitoes collected in the field were at immature stages, and they were reared to adults under laboratory conditions at 25±4°C and a 12:12 h photoperiod. They were morphologically identified and stored at −20°C until DNA extraction.

### DNA isolation

DNA was isolated from ∼30 individual mosquitoes per location using the salt extraction technique [23] and resuspended in 50 μL of water (Cellgro® sterile purified water, molecular biology grade and free of proteases, DNases and RNases). The concentration and quality of each DNA sample was determined on a ThermoScientific NanoDrop 2000 spectrophotometer.

### Development of the multiplex PCR method

The amplification primers used for the variants of loci 410, 1,016 and 1,534 are given in Table 1.

**Table 1.**
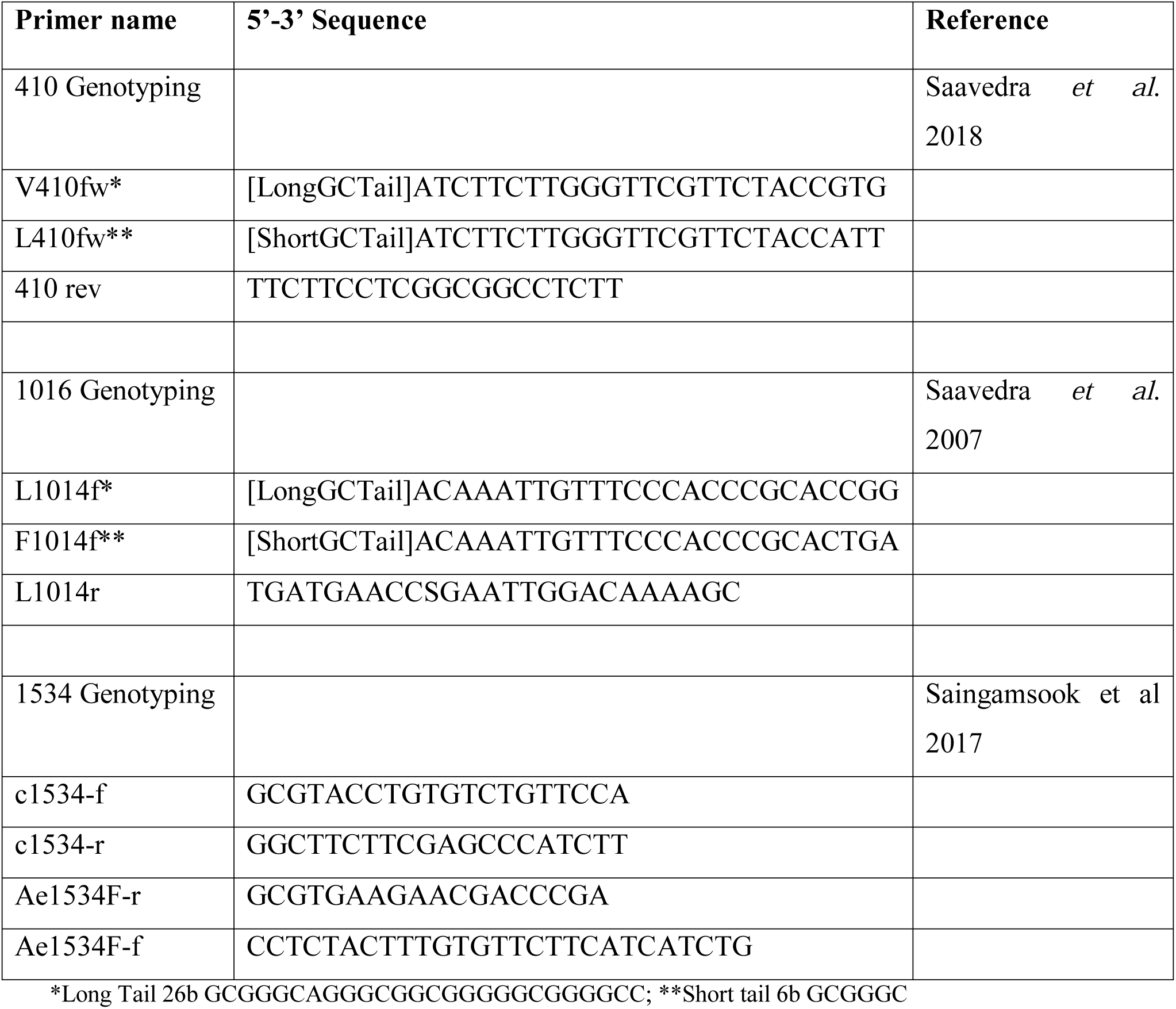
Sequences of the primers used for multiplex PCR.

The specific oligonucleotides V410fw and L410fw amplify a region of 113 bp, corresponding to the L410 allele (resistant). The specific oligonucleotides V410fw and 410rev amplify a region of 133 bp, corresponding to the V410 allele (susceptible). The specific oligonucleotides I1,016f and I1,016r amplify a region of 82 bp, corresponding to the I1,016 allele (resistant). Oligonucleotides V1,016f and I1,016r amplify a region of 102 bp, corresponding to the V1,016 allele (susceptible). The oligonucleotides c1,534-f and c1,534-r amplify a region of 368 bp with which the specific oligonucleotides Ae1,534F-r and Ae1,534C-f hybridize, resulting in products of 180 bp for the C1,534 allele (resistant) and of 232 bp for the F1,534 allele (susceptible) (Figure 1).

**Fig. 1.**
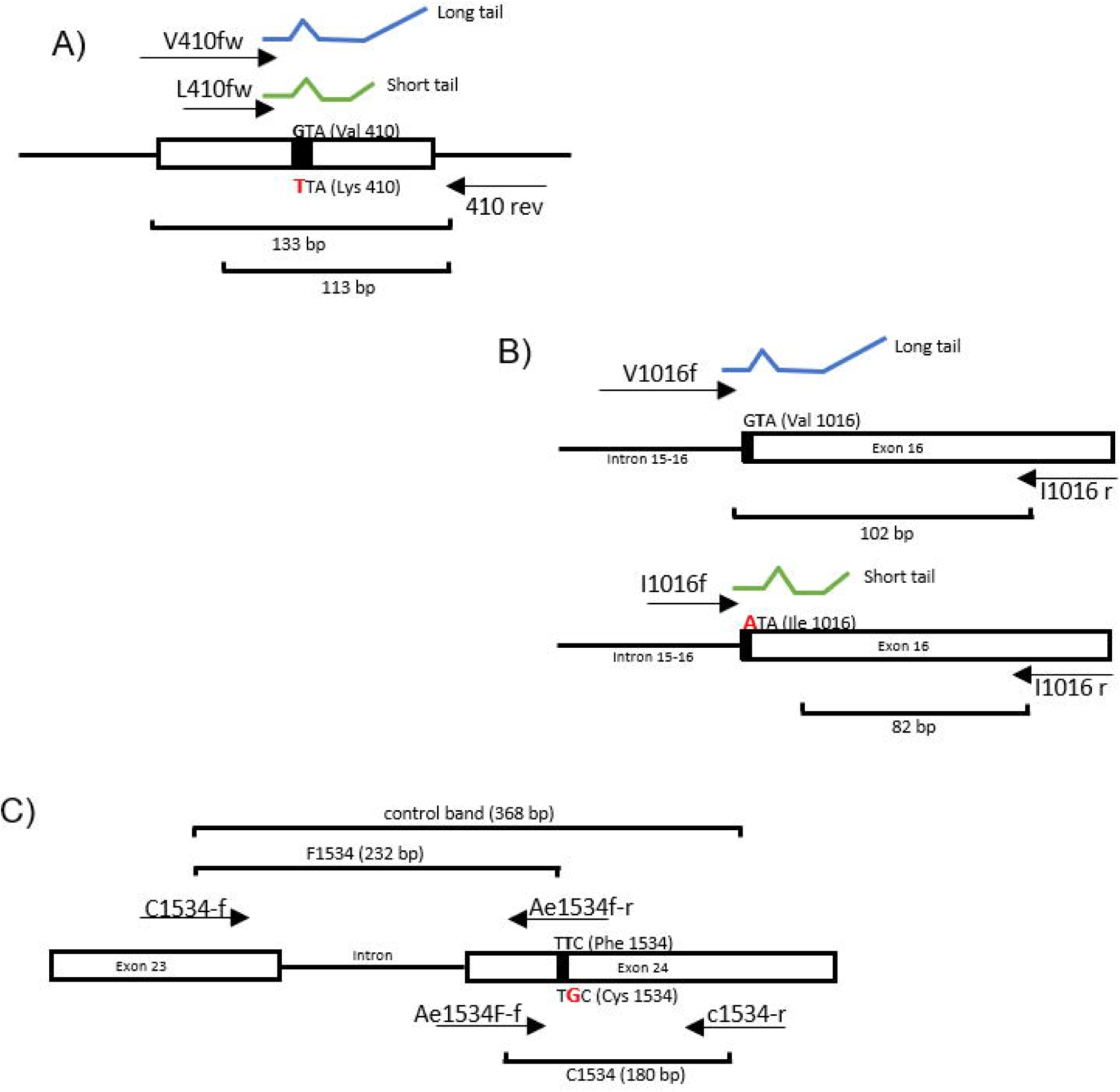
Schematic of the AS-PCR assay for detection of the A) V410L, B) V1,016I and C) F1,534C mutations.

Tests for optimization of PCR conditions resulted in the following multiplex PCR protocol. The DNA samples used in the amplification process were in a concentration range of 20-250 ng/μL. The final reaction mixture was 19.12 μL and contained: 1.02X buffer, 1.53 mM MgCl_2_, 0.2 mM dNTPs, oligonucleotides for genotyping 410 at a final reaction concentration of 1.27 pmol/μL, for 1,016 at 1.02 pmol/μL and for 1,534 at 0.82 pmol/μL (Table 1), and also 5 U Taq DNA polymerase.

The reaction was carried out in a Multigene Optimax thermal cycler (Labnet International, Edison, NJ, USA). The reaction conditions were as follows: 95°C for 2 min for the initial separation of the DNA strands, followed by 45 cycles of 95°C (30 s), 58.6°C (1 min) and 72°C (30 s) and a final extension for 2 min.

A PCR tube containing all the components except genomic DNA was run with the primers as a contamination control. Controls were included in each PCR performed, the New Orleans strains was used as susceptible control.

After amplification, 4 µl of the products of the PCR reaction mixture were analyzed by horizontal electrophoresis on a 2.5% agarose gel. The electrophoresis conditions were 110 V for 1 h using 1X SB buffer (200 mM sodium borate buffer, pH 8) along with a 25-bp molecular weight marker to determine the size of the fragments and staining with GelRed® (Biotium, Hayward CA, USA). The PCR products were visualized with a transilluminator (UVITEC, Cambridge, UK). At the end of amplification, it was possible to obtain up to 7 PCR products per sample, whose size indicated the genotypic combination for loci 410, 1,016 and 1,534.

### Validation of the study

For the validation of the results obtained in multiplex PCR, AS-PCR was performed according to Saavedra et al. [10], Yanola et al. [24] and Villanueva-Segura et al. [15].

## Results

### Multiplex PCR vs AS-PCR assay

Mosquito DNA of each population was used to detect the mutations V410L, V1,016I and F1,534C. Molecular assays were conducted on 147 mosquitoes of the populations analyzed. The genotype L410/L410 (homozygous mutant) was seen as a single band of 113 bp and the homozygous wild-type genotype (V410/V410) as a single band of 133 bp, while the heterozygous genotype (V410/L410) showed both bands. Genotype I1,016/I1,016 (homozygous mutant) was seen as a single band of 82bp and the homozygous wild-type genotype (V1,016/V1,016) as a single band of 102 bp, while the heterozygous genotype (V1,016/I1,016) showed both bands. The homozygous wild-type genotype (F1,534/F1,534) showed a single band of 232 bp and the homozygous mutant genotype (C1,534/C1,534) a single band of 180 bp, while the heterozygous genotype (F1,534/C1,534) had both bands.

In the case of the detection of the F1534C mutation, an internal control band of 368 bp is obtained (Figure 2).

**Fig. 2.**
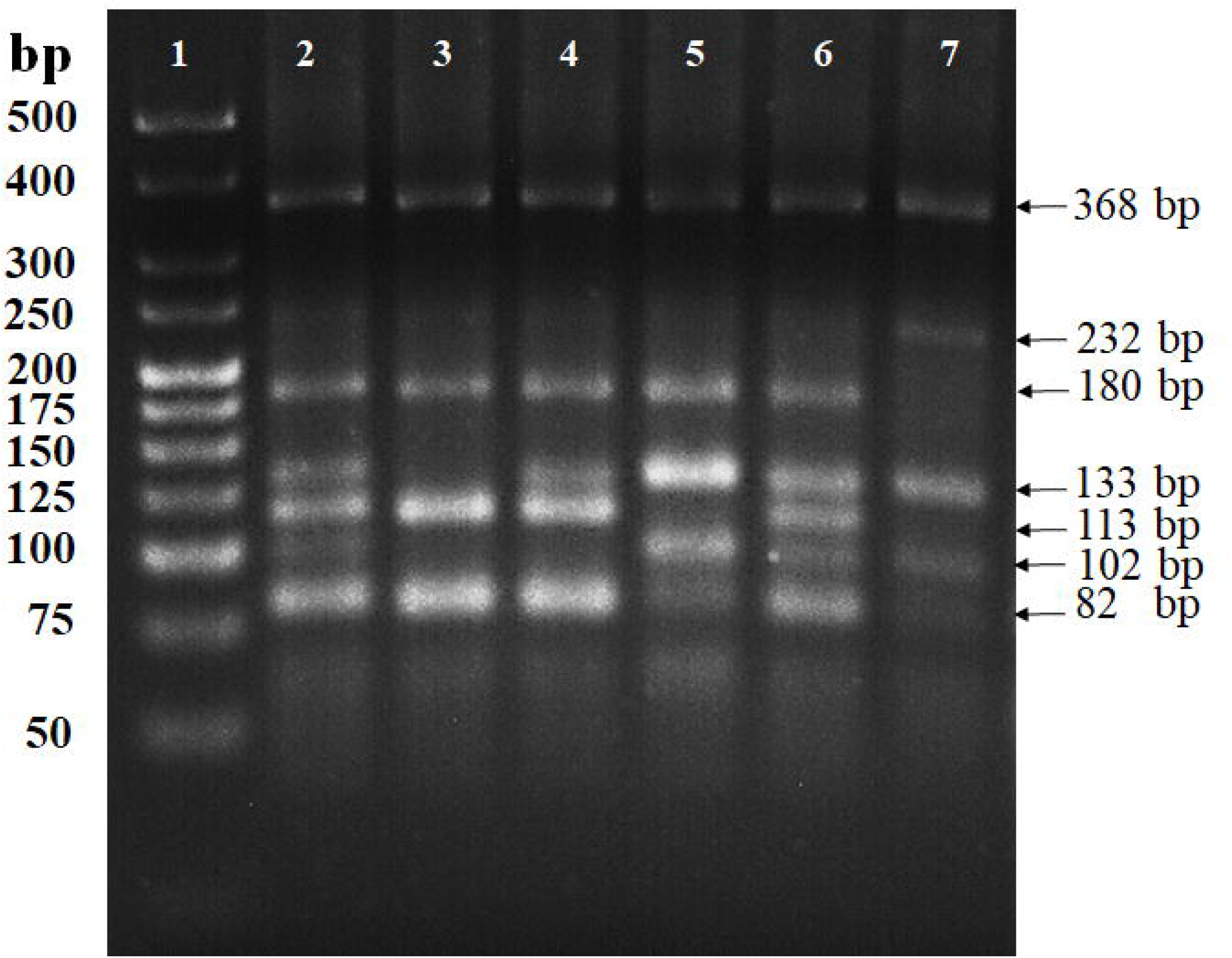
Agarose gel 2.5% electrophoresis of amplified gDNA products in several mosquitoes using multiplex PCR. The arrows indicate a common band of 368 bp, a band of 232 bp corresponds to the susceptible allele F1,534, a band of 180 bp corresponds to the resistant allele C1,534, a band of 133 bp corresponds to the susceptible allele V410, a band of 113 bp corresponds to the resistant allele L410, a band of 102 bp corresponds to the susceptible allele V1,016, a band of 82 bp corresponds to the resistant allele I1,016. Lane 1 shows the molecular weight marker (25pb DNA Ladder). Lane 2 corresponds to a resistant homozygous (C1,534 / C1,534), heterozygous (V410 / L410) and heterozygous (V1,016 / I1,016) individual. Lane 3 shows a resistant homozygous individual (C1,534 / C1,534), resistant homozygous (L410 / L410) and resistant homozygous (I1016 / I1016). Lane 4 corresponds to a resistant homozygous individual (C1,534 / C1,534), heterozygous (V410 / L410) and resistant homozygous (I1,016 / I1,016). Lane 5 corresponds to a resistant homozygous individual (C1,534 / C1,534), susceptible homozygous (V410 / V410) and heterozygous (V1,016 / I1,016). Lane 6 corresponds to a resistant homozygous individual (C1,534 / C1,534), heterozygous (V410 / L410) and heterozygous (V1,016 / I1,016). Lane 7 corresponds to a triple susceptible homozygous individual (V410 / V410, V1,016 / V1,016, F1,534 / F1,534).

A total of 12 genotypes were obtained and validated in the populations analyzed, using multiplex PCR and the individual endpoint technique previously reported by Saavedra et al. [10], Yanola et al. [24] and Villanueva-Segura *et al*. [15], demonstrating that all samples had the same genotype using multiplex PCR as with AS-PCR (Tables 2 and 3).

**Table 2.**
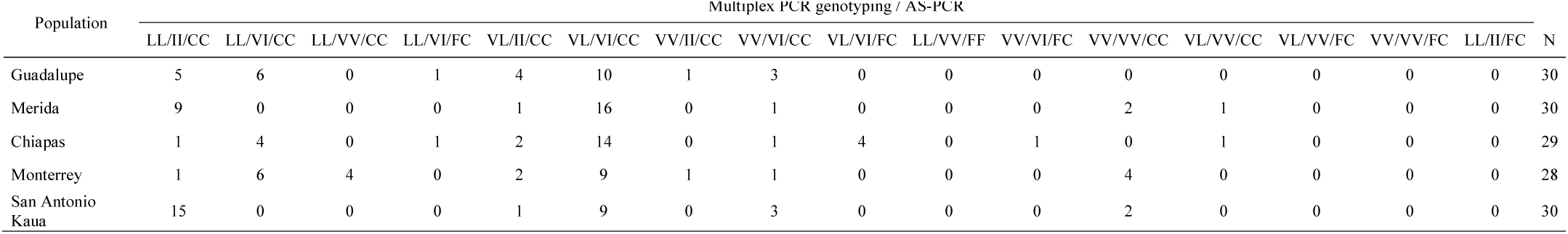
Comparison of genotyping results for V410, V1016, F1534 mutations from multiplex PCR and AS-PCR.

**Table 3.**
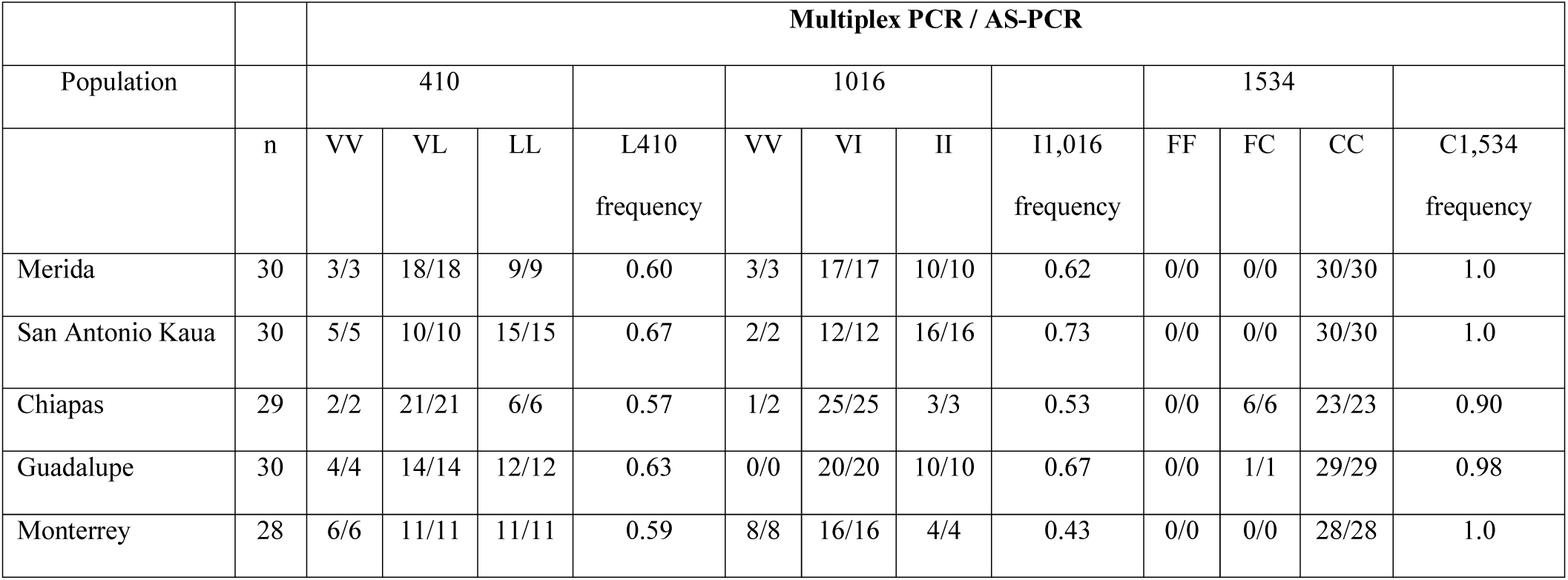
Comparison of genotypes for V410L, V1016I, and F1534C mutations for multiplex PCR and AS-PCR.

|  |  | **Multiplex PCR / AS-PCR** |  |  |  |  |  |  |  |  |  |  |  |
| --- | --- | --- | --- | --- | --- | --- | --- | --- | --- | --- | --- | --- | --- |
| Population |  | 410 |  |  |  | 1016 |  |  |  | 1534 |  |  |  |
|  | n | VV | VL | LL | L410<br>frequency | VV | VI | II | I1,016<br>frequency | FF | FC | CC | C1,534<br>frequency |
| Merida | 30 | 3/3 | 18/18 | 9/9 | 0.60 | 3/3 | 17/17 | 10/10 | 0.62 | 0/0 | 0/0 | 30/30 | 1.0 |
| San Antonio Kaua | 30 | 5/5 | 10/10 | 15/15 | 0.67 | 2/2 | 12/12 | 16/16 | 0.73 | 0/0 | 0/0 | 30/30 | 1.0 |
| Chiapas | 29 | 2/2 | 21/21 | 6/6 | 0.57 | 1/2 | 25/25 | 3/3 | 0.53 | 0/0 | 6/6 | 23/23 | 0.90 |
| Guadalupe | 30 | 4/4 | 14/14 | 12/12 | 0.63 | 0/0 | 20/20 | 10/10 | 0.67 | 0/0 | 1/1 | 29/29 | 0.98 |
| Monterrey | 28 | 6/6 | 11/11 | 11/11 | 0.59 | 8/8 | 16/16 | 4/4 | 0.43 | 0/0 | 0/0 | 28/28 | 1.0 |

The analysis of the frequencies of the L410, I1,016 and C1,534 was determined in the 147 mosquitoes of the five selected populations. The frequencies for the allele L410 in the population of Chiapas was 0.57, in Merida 0.60 and in San Antonio Kaua 0.67, while the frequencies were 0.63 and 0.59 for the populations of Guadalupe and Monterrey, respectively. The frequencies for I1,016 in the populations of Merida, San Antonio Kaua, Chiapas, Guadalupe and Monterrey were 0.62, 0.73, 0.53, 0.67 and 0.43, respectively, and the mutation F1534C was fixed in the populations of Merida, San Antonio Kaua and Monterrey and with frequencies of 0.90 and 0.98 for the populations of Chiapas and Guadalupe, respectively (Table 3).

In the populations analyzed, co-occurrence of homozygous resistant genotypes was greatest in the population of San Antonio Kaua, followed by the population of Merida, both from the state of Yucatan. The lowest levels of triple homozygous resistant were in the populations of Monterrey (0.04) and Chiapas (0.03). The most frequent genotype was double heterozygous for 410 and 1,016 and homozygous resistant for 1,534 (V410/L410, V1,016/I1,016, C1,534/C1,534) (Table 4).

**Table 4.** Frequency of co-occurrence of the mutations V1016I, F1534C y V410L in populations of *Ae. aegypti*

| Population | Genotype frequency V1016I, F1534C y V410L |  |  |  |  |  |  |  |  |  |  |  |
| --- | --- | --- | --- | --- | --- | --- | --- | --- | --- | --- | --- | --- |
|  | LL/II/CC | LL/VI/CC | LL/VV/CC | LL/VI/FC | VL/II/CC | VL/VI/CC | VV/II/CC | VV/VI/CC | VL/VI/FC | VV/VI/FC | VV/VV/CC | VL/VV/CC |
| Merida | 0.30 | 0.00 | 0.00 | 0.00 | 0.03 | 0.53 | 0.00 | 0.03 | 0.00 | 0.00 | 0.07 | 0.03 |
| San Antonio Kaua | 0.50 | 0.00 | 0.00 | 0.00 | 0.03 | 0.30 | 0.00 | 0.10 | 0.00 | 0.00 | 0.07 | 0.00 |
| Chiapas | 0.03 | 0.14 | 0.00 | 0.03 | 0.07 | 0.48 | 0.00 | 0.03 | 0.14 | 0.03 | 0.00 | 0.03 |
| Guadalupe | 0.17 | 0.20 | 0.00 | 0.03 | 0.13 | 0.33 | 0.03 | 0.10 | 0.00 | 0.00 | 0.00 | 0.00 |
| Monterrey | 0.04 | 0.21 | 0.14 | 0.00 | 0.07 | 0.32 | 0.04 | 0.04 | 0.00 | 0.00 | 0.14 | 0.00 |

## Discussion

Performing PCR even today still takes time and effort; however, the objective of developing a multiplex PCR method is mainly to reduce these factors by being able to amplify various alleles in the same reaction. This is difficult when it is time to standardize the method, where each pair of primers included increases the difficulty since it is not enough to match the Tm (melting temperature) and AG (adenosine:guanidine) of these, so extra effort is needed in their design [25].

The optimization of every multiplex PCR method has critical difficulties. The design of the primers is key to a successful PCR, and the presence of more than one pair increases the possibility of dimers and also requires the adjustment of the other PCR components (buffers, dNTPs, MgCl_2_ and Taq DNA polymerase)[26].

One of the main utilities of multiplex PCR is the simultaneous detection of multiple genes, such as serogroups of pathogens like with *Salmonella, Escherichia coli* O157 and *Listeria monocytogenes* [27, 28], including allele-specific multiplex PCR for detection of multiresistance in *Mycobacterium* tuberculosis [29]. However, the utility of multiplex PCR is not limited to this; one of the first reports of detection of multiple kdr mutations for a single L1014L/S site by multiplex PCR was performed by Tan *et al*. [30] for the alleles L/S (TTG/TCG), L/F (TTG/TTT) and L/L (TTG/TTG). Assays have been performed for the detection of V1,016I and F1,534C mutations individually [10, 24], and as multiplex PCR with other mutations such as V1,016G and F1,534C, T1520I and F1,534C [21, 31]. The V410L mutation was first detected in co-occurrence with the V1,016I and F1,534C mutations in *Ae. aegypti* mosquitoes but only by simplex PCR [32].

The utility of multiplex PCR is also not limited only to the detection of a single resistance mechanism. Kazanidou *et al*. [33] standardized the detection of polymorphisms by the substitution of five nucleotides in the sodium channel gene and the ace-1 gene ([kdr-w homozygous], [kdr-e homozygous], [kdr heterozygous], ace-1r homozygous, and their hybrids [ace1s/ace-1r, kdrs/kdr-w]) in only one of the samples.

In this study, we used a simplex PCR as a confirmation technique to characterize each mutation in the samples used and to validate the multiplex PCR method. Six samples were used among which we could determine up to 27 different genotypes. The results of the test validation showed 100% agreement between multiplex PCR and simplex PCR. These results indicated high sensitivity and specificity values for the new multiplex PCR method developed. This method allows us to determine the frequency of multiple mutations, as well as their co-occurrence, in a single reaction [28].

Our method is the first designed for multiplex genotyping of the three kdr mutations in *Ae. aegypti* in Mexico, i.e., V410L, V1,016I, and F1,534C.

## Conclusion

The multiplex PCR technique allows simultaneous and reliable detection of the V410L, V1,016I and F1,534C mutations in *Ae. aegypti* populations in Mexico. This method optimizes the monitoring times of mutant alleles and genotypic frequencies in wild populations of this species. The characteristics of this technique make it an advantageous alternative for countries where the dynamics of pyrethroid resistance in mosquitoes is variable and changing and where resources for this purpose are also limited.

## Declarations

## Acknowledgments

Dr. A. Leyva (USA) provided English translation and editing of the manuscript.

## Availability of data and materials

All data generated or analyzed during this study are included in this published article. The raw datasets generated and /or analyzed during the current study are available from the corresponding author upon reasonable request.

## Consent for publication

Not applicable

## Authors’ contributions

AEF conceived the study and funds acquisition; OKVS coordinated the experiments; GPG, EdJMJ, KOZ, FJGS carried out collections of the populations included in the study; LP, EdJMJ, FJGS, KOZ performed the lab work; AEF, JADB, BLM analyzed data; AEF, OKVS, JADB drafted the manuscript; JADB, BLM, GPG provided critical input regarding the findings. All authors read and approved the final version of the manuscript.

